# Comprehensive Characterization of the Human Neural Stem Cell Line HNSC.100 as a Versatile Model for Neurobiological Research

**DOI:** 10.64898/2026.01.21.700829

**Authors:** Estera Jeruzalska, Carolin Ketteler, Elisa Stützenberger, Sandra Burczyk, Larissa Möller, Dierk Niessing

**Author notes:** These authors contributed equally (Estera Jeruzalska, Carolin Ketteler).

## Abstract

Studying neural-related questions is inherently challenging due to the limited number of suitable cell models. Here, we characterize a previously reported immortalized human neural stem cell line, HNSC.100, serving as a robust model for a wide range of neurobiological research questions. The cell line expresses key neural stem cell markers, including SOX2, vimentin, nestin, and allows for efficient genetic manipulation. Furthermore, HNSC.100 cells can be differentiated into neurons, astrocytes, and oligodendrocytes, thereby covering a wide spectrum of major neural cell types. We adapted corresponding differentiation protocols and established a comprehensive panel of molecular markers to validate successful differentiation, enabling precise characterization of the resulting cell population. In addition, we provide a complete dataset of RNA expression levels for all detectable genes in HSNC.100 cells. Based on this dataset, we assembled a list of expressed genes implicated in neural disorders that can be studied with this cell line. Together, we present a detailed characterization of the HNSC.100 cell line and provide new tools and reference data to facilitate its use. This resource enables researchers to evaluate the line’s suitability for specific applications and to rapidly integrate HNSC.100 cells into their experimental workflows.

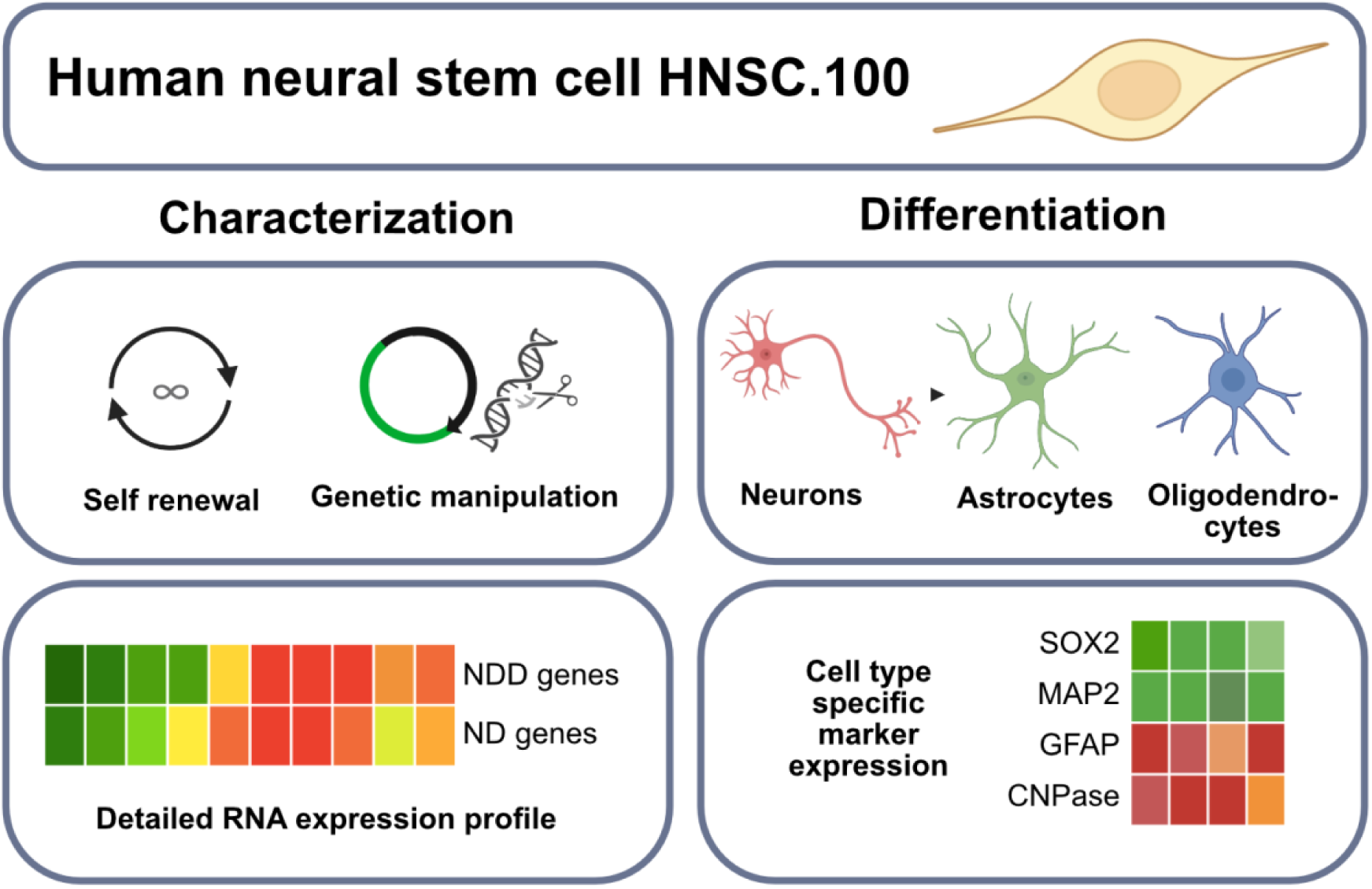

## Introduction

Biological processes and their underlying molecular functions can be studied in many ways, for instance by using *in vitro-* and cell culture-based approaches or various model organisms. Cultures of immortalized human cells have been the workhorse for decades, allowing us to study fundamental biological processes. The ease of handling, high efficiency of genomic modifications, and their replication time render them the first choice for many research projects. However, when using immortalized cells, it is also important to consider their downsides, including unphysiological properties due to their carcinogenic origin or immortalization. These features frequently result in genomic abnormalities and instability, as well as limited differentiation plasticity. Such cell lines include Jurkat, HeLa, HEK293, K562, and MCF-7 cultures, to name a few. While they are great models to study certain cellular functions, they might not be applicable for addressing other biological questions. Such limitations become most apparent when studying cell types such as neural stem cells that have great differentiation potential under physiological conditions. Moreover, when investigating neuronal disorders, the gene expression profiles of the above-mentioned commonly used immortalized cell lines differ significantly from those of neural and neuronal cells. Certain transcripts and proteins are specific to a particular cell type and may be entirely absent in others.

To date, the number of available immortalized neur(on)al cell lines is limited, and for existing models it is often difficult to balance between the ease of use and their potential to act as functional neur(on)al cells. This challenge becomes most obvious when considering that cell cultures rely on their proliferative capacity, whereas neuronal or glial cells are rather postmitotic. Although both iPSC-derived neural stem cells (NSCs) and rodent primary neuronal cultures are considered more physiological than immortalized cells, each presents distinct limitations. iPSC-derived NSCs, generated, for instance, from reprogrammed human fibroblasts [1,2], retain a diploid human genetic background and can be patient-specific. However, time-consuming and long differentiation protocols carry the risk of high variability between different NSC lines. Additionally, they typically have a limited expansion window, as prolonged passaging can lead to spontaneous differentiation or loss of NSC identity. In contrast, primary rodent neurons [3,4] provide direct access to postmitotic neuronal cells, but they are limited by low yield, short lifespan *in vitro*, and species-specific differences that limit translational relevance. Importantly, due to their limited proliferation and scalability, iPSC-derived as well as rodent primary neural cell types are poorly suited for techniques requiring large cell numbers, such as proteomics, metabolomics, CLIP, or BioID. In contrast, immortalized neural cell lines offer easier handling, high proliferation rates, while maintaining stem cell characteristics and greater reproducibility between different studies. These features render them especially well-suited for high-throughput studies.

Amongst the most widely used immortalized human cells with neural properties and differentiation potential are SK-N-SH and the SH-SY5Y cells [5–7], as well as the cell lines ReNcell CX and VM [8]. SK-N-SH cells were derived in 1973 from a neuroblastoma patient [9] and were established as a heterogeneous mixture of cells. From those cells, the monoclonal SH-SY5Y cell line was derived. SH-SY5Y cells are widely used due to their ease of cultivation and because of their potential to differentiate into cells that form neurite networks upon stimulation with retinoic acid or phorbol esters. Other differentiation protocols have been explored that can drive differentiation into different subtypes of neurons [10]. However, a major limitation when studying basic features of neural cells is their inability to differentiate into cell types other than neurons. In contrast to SH-SY5Y cells, cell lines such as ReNcell VM and CX have the capacity to differentiate into neurons, astrocytes, and oligodendrocytes under specific conditions [8,11]. These human neural progenitor cell lines were derived from ventral mesencephalic (VM) and cortical (CX) regions of human fetal brains. Their immortalization using v-myc (VM) or c-myc (CX) oncogenes allowed for long-term proliferation while retaining their differentiation capacity. Given the very limited number of similar cell lines, the availability of other immortalized neural cell lines would be beneficial, for instance, to allow for comparative studies and thus reducing the risk of cell type-specific artefacts.

Here we describe the human non-cancerous neural stem-cell line HNSC.100, also referred to as hNS1, as an alternative model to the above-mentioned cell types. HNSC.100 cells were derived from the forebrain of an aborted fetus and immortalized using *retroviral transduction of p110 gag-myc* gene [12]. The established cell line is nestin-positive and self-renewing. The successful integration of the retroviral gene was confirmed using the Southern Blot, and the proliferation capacity was confirmed using the [^3^H] Thymidine Incorporation assay.

The original publication from 2000 already reported that these cells express lineage-specific markers upon withdrawal of growth factors, indicative of differentiation into neurons (MAP2, β-tubulin), astrocytes (GFAP), and oligodendrocytes (GalC). With this feature, they exhibit a much greater diversity in differentiation potential than, for instance, SH-SY5Y cells. Consistently, studies using HNSC.100 cells have focused on neuronal differentiation [13] and neurodegeneration-relevant applications, including Parkinson’s disease–related dopaminergic differentiation paradigms [14] and transplantation studies in the adult rodent brain [15]. Despite these advantages and the need for additional cell lines with neural differentiation potential, HNSC.100 cultures are not yet widely used. A major reason for this might be the lack of a detailed characterization of the HNSC.100 cell line and the thus-far limited number of reported standardized tools.

To overcome these limitations, we provide a detailed description of the HNSC.100 cell line, improved differentiation protocols, and established a toolbox of validated primers and antibodies to track differentiation states by real-time quantitative PCR (qPCR) and immunofluorescence. Furthermore, to allow for a fast assessment of whether HNSC.100 cells are suitable for a particular project, we performed a karyotype analysis and provided an RNAseq-based list of genes with reasonable expression levels. We believe that the information provided here will allow for a quick assessment if the HNSC.100 cells are suitable for a given project, and the rapid establishment of this cell line in standard cell-biology laboratories.

## Results

### Cultivation of HNSC.100 cells

As described earlier, the HNSC.100 cells were cultured on poly-D-lysin-coated surfaces in a defined medium supplemented with the mitogenic factors EGF and FGF-2 [12]. Under these conditions, the cells demonstrated robust proliferative capacity across multiple passages. Like other immortalized cells, HNSC.100 cells are prone to karyotypic abnormalities, including multiple regional copy number gains and losses (Supplementary Figure 1; Supplementary Table 1). However, these genomic abnormalities seem to have no noticeable impact on cell survival, proliferation, or differentiation capacity. Growth kinetics showed a characteristic exponential growth curve with a doubling time of 1.72 days (Fig. 1B, C), consistent with the original publication (1.7 +/- 0.08 days) [12]. The HNSC.100 cells grew as a uniform monolayer and had a neural-like bipolar shape, characterized by the presence of small neurite-like protrusions (Fig. 1B).

**Fig. 1.**
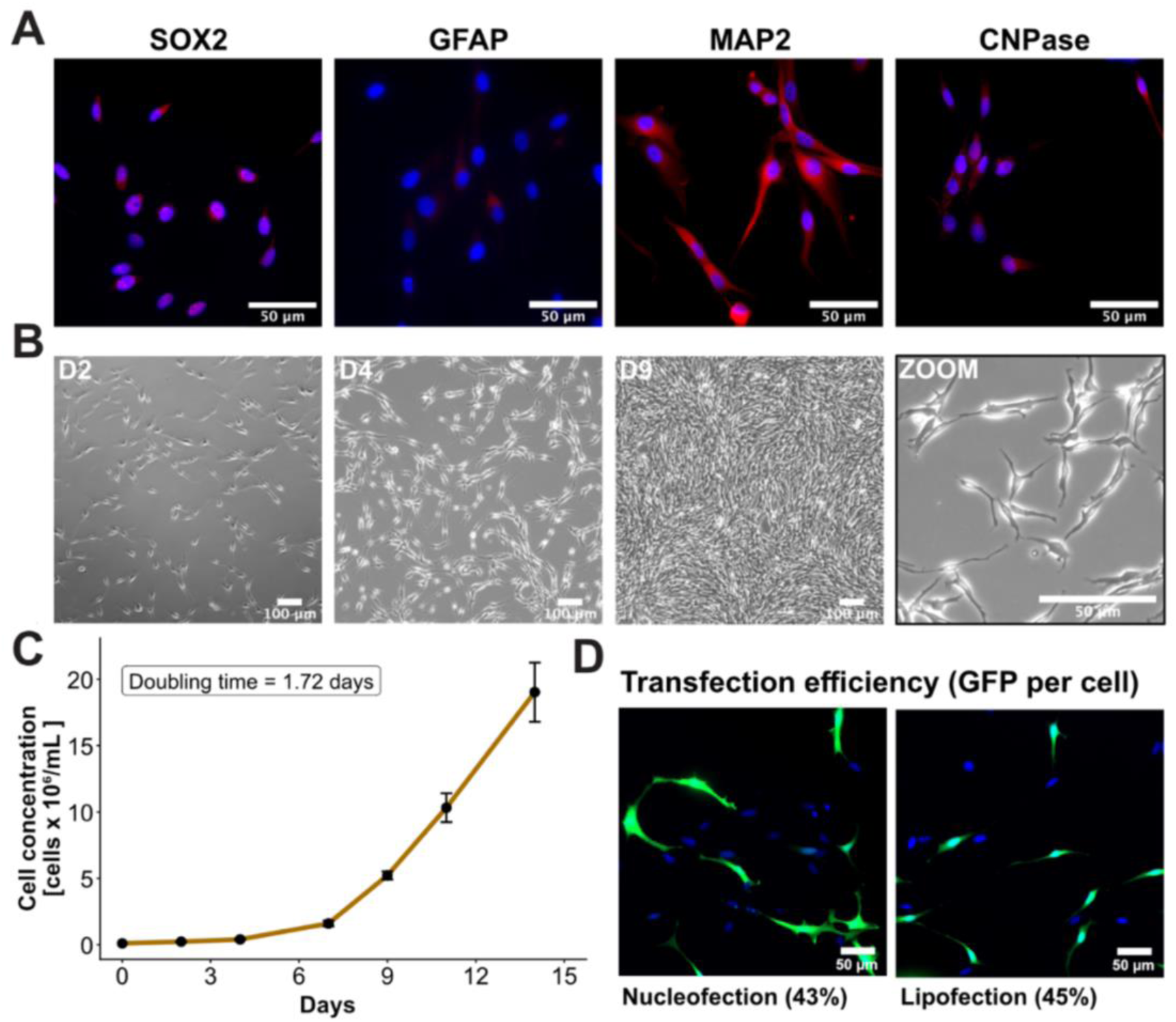
General characteristics of the HNSC.100 cell line. **A.** Immunofluorescence staining of cell type-specific markers revealed that every HNSC.100 cell is SOX2-positive, indicating their multipotent state. Additionally, cells show no or weak expression of GFAP and CNPase, respectively, but a variable expression of MAP2. **B.** Growth of cells within a week after seeding. After about seven days, confluency of cells in the flask is reached, indicating the need to passage them. The zoomed-in image shows the neural-like shape of HNSC.100 cells from day 4. **C.** Quantification of cell growth over time. Cells were cultivated in triplicates, every 2-3 days detached, counted, and reseeded. Based on these measurements, a duplication rate of 1.72 per day was calculated. **D)** Transfection efficiency was determined using the Countess III FL (Thermo Fisher) and immunofluorescence pictures from both approaches are shown.

Immunofluorescence analysis revealed a strong expression of the neural stem cell marker SOX2, confirming their undifferentiated multipotent state (Fig. 1A and Supplementary Figure 2) [16,17]. To obtain a broader picture of the state of the HNSC.100 line, we performed bulk RNA sequencing and observed that neural stem cell markers, such as nestin and vimentin, were also highly expressed at the RNA level (Supplementary Table 3). In contrast, markers for astrocytes and oligodendrocytes, like GFAP and CNPase, were expressed at low RNA levels and barely detected in the immunofluorescence images (Fig. 1A and Supplementary Table 3), further confirming the undifferentiated state of the cells. Moreover, the neuronal marker MAP2 was detected on RNA and protein levels in HNSC.100 cells, although neuronal differentiation has not been initiated. Heatmap analysis of the bulk RNA sequencing data confirmed high expression of neural stem cell and proliferation markers (SOX2, NES, VIM, MKI67) and comparatively low expression of neuronal, astrocytic, and oligodendrocytic differentiation markers (Supplementary Figure 3). This pattern was also observed when comparing the HNSC.100 line with publicly available RNA-sequencing datasets from other immortalized human neural stem cells (ReNcell CX and VM) [18,19] as well as iPSC-derived neural progenitor cells (NPCs) [20] (Supplementary Figure 3). Notably, NSC and proliferation markers showed the highest expression when compared to differentiation markers in the iPSC-derived NPCs and the HNSC.100 cell line. In contrast, the ReNcell CX and ReNcell VM showed a less distinct separation, with some differentiation markers reaching expression levels exceeding NSC and proliferation markers. This observation suggests that HNSC.100 provides a useful expansion of the spectrum of immortalized neural stem cell lines and offers additional opportunities for orthogonal studies.

Together, these data characterize the HNSC.100 as an immortalized, SOX2-positive, highly proliferating human neural stem cell line maintained under well-defined conditions.

### Cell and genome manipulation

For a detailed characterization of the biological functions of a given gene and its protein product, various methods for gene manipulation are available. These include gene depletion by shRNA or CRISPR/Cas9 technologies, and overexpression, possibly with added gene modifications, via transient or stable transfection, potentially followed by single-cell cloning. In particular for selection-based approaches and subsequent high-throughput techniques, easy-to-handle, stable, immortalized cells offer great advantages. We have successfully performed various genetic manipulations in the HNSC.100 cell line using lipofection or nucleofection methods, including knock-down and CRISPR/Cas-based knock-out of genes, as well as the generation of transiently and stably expressing cell lines. Using a GFP-expressing control vector (pmaxGFP^TM^, Lonza) with the 4D-Nucleofector (Lonza) and the Lipofectamine 3000 transfection kit (L3000, Invitrogen™), we determined transfection efficiency to be in the range of 43-45% (Fig. 1D). Depending on the application, a homogenous HNSC.100 line derived from a single cell may be obtained in about 6-8 weeks.

### Neural differentiation

The first study on HNSC.100 cells included a description of their spontaneous differentiation into astrocytes, neurons, and oligodendrocytes upon removal of growth factors. As a result, a combination of GFAP-, MAP2-, and TUBB3- positive cells plus a small fraction of Gal-C-positive cells was obtained [12]. In our hands, this original procedure primarily leads to the differentiation into astrocytes and not into neurons or oligodendrocytes. For this astrocyte-specific protocol, we maintained the cells for up to 16 days until they no longer showed morphological changes (Fig. 2; Supplementary Table 2). Notably, at this stage cells exhibited a typical star-like morphology with their soma surrounded by numerous branches [21,22]. However, we also noticed a few bipolar cells that appeared more characteristic of neuronal cells (Fig. 2). Since the original protocol predominantly yielded astrocyte-like cells, alternative protocols were explored to promote differentiation into other lineages. First, we tested protocols by the company Thermo Fisher for the differentiation of HNSC.100 into neurons and oligodendrocytes [23]. Their neuronal differentiation protocol yielded a heterogeneous mixture of neurons with uni-, bi-, and multi-polar cells after 28 days (Fig. 2; Supplementary Table 2), a time point where cell morphologies had stopped changing. These neurons exhibited very thin and long neurite outgrowths consistent with typical neuronal characteristics. On the other hand, the recommended conditions for oligodendrocyte differentiation did not yield successful results in our hands, prompting us to explore alternative strategies. Finally, adapting a two-step cultivation procedure originally established for human embryonic stem cells led to successful oligodendrocyte differentiation (Supplementary Table 2) [24]. This procedure lasted for 35 days and resulted in a mixture of small and round cell bodies surrounded by many branches, and more polygonal-shaped cell bodies with fewer observed processes (Fig. 2). This suggests the presence of both type I and II oligodendrocytes [25,26].

**Fig. 2.**
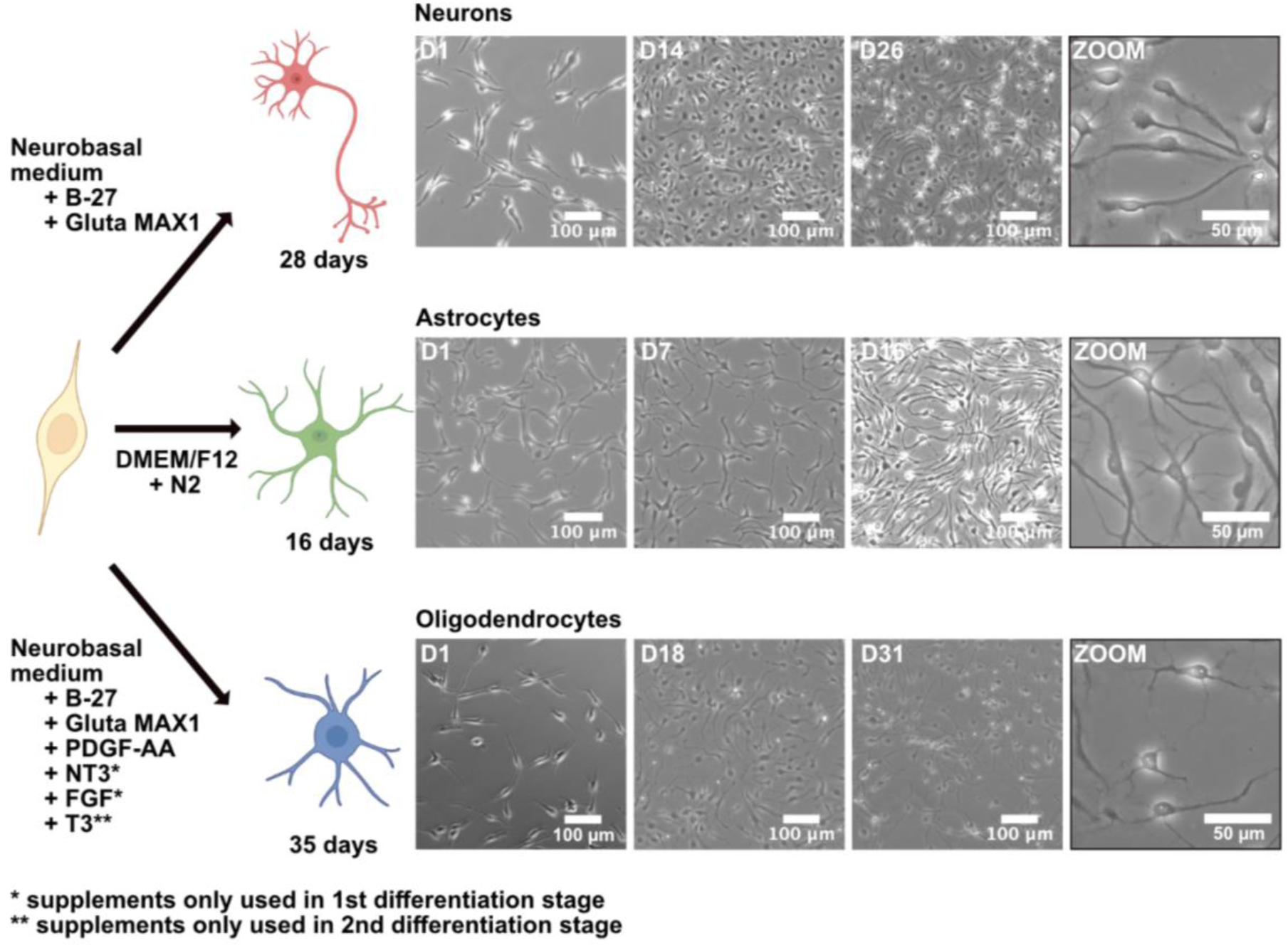
Cell type-directed differentiation of HNSC.100 line. Overview of key medium compositions for the generation of specific cell types, and respective time frames required for differentiation. For oligodendrocytes, two distinct steps were necessary, indicated as 1^st^ and 2^nd^ differentiation stages (marked by * and **, respectively). For a detailed description of medium compositions and protocols see Supplementary Table 2 and Materials & Methods. Representative images show the morphological changes during differentiation into neurons (upper panel), astrocytes (middle panel), and oligodendrocytes (lower panel). Representative zoom-in images were taken from the last day of differentiation.

To complement our morphology-based cell-type analysis with molecular features, we also assessed the different cell populations with cell-type-specific markers. We first used some of the most frequently employed differentiation markers, including SOX2, GFAP, MAP2, and CNPase [27,28], and performed transcript quantification via qPCR and protein expression analysis via immunofluorescence staining (Fig. 3). The analysis showed that undifferentiated HNSC.100, but none of the differentiated cells were positive for the neural stem cell marker SOX2 on the protein level (Fig. 1A & 3B and Supplementary Figure 2), although they exhibited higher expression on the transcript level (Fig. 3A).

**Fig. 3.**
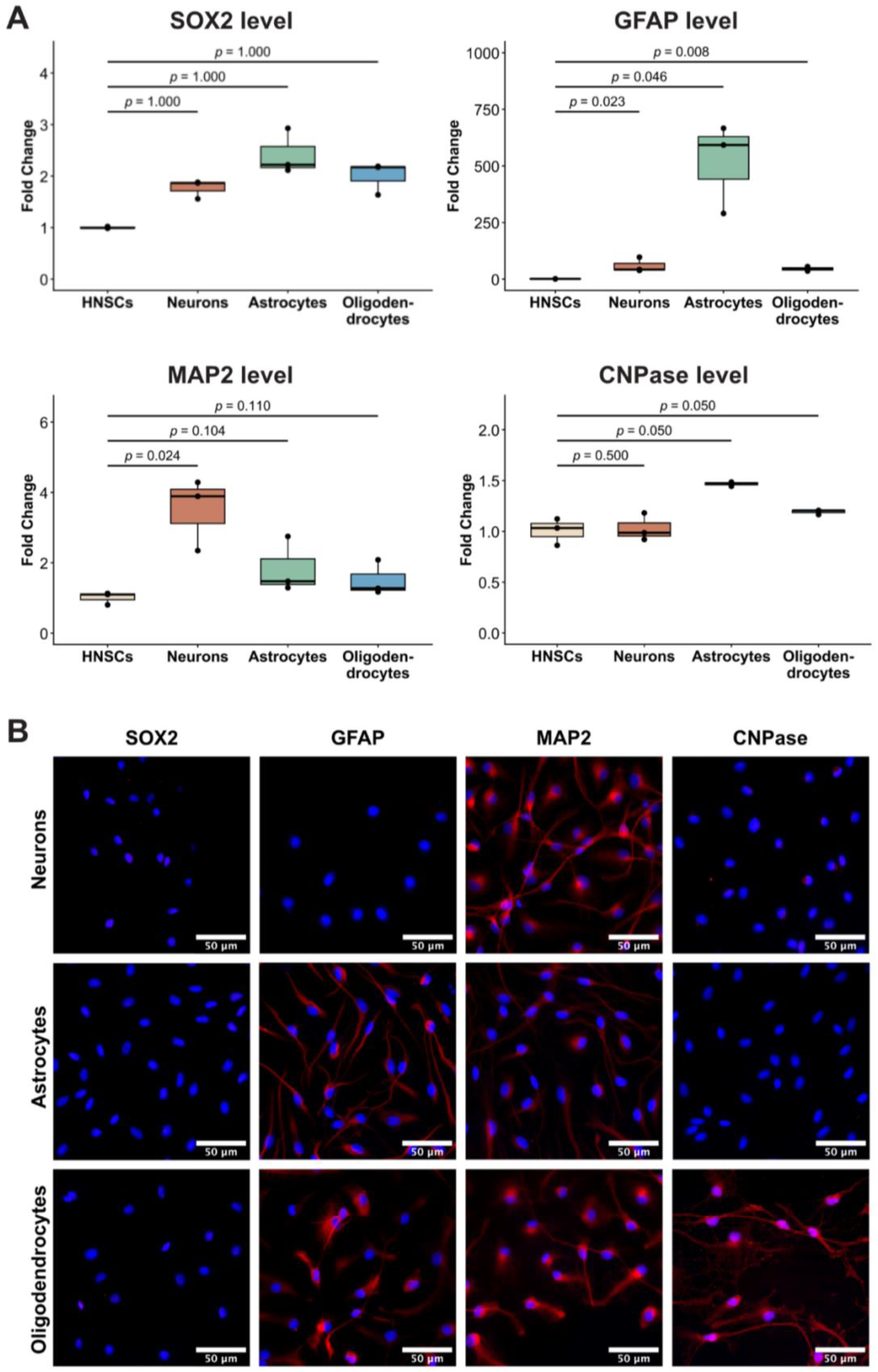
Validation of cell differentiation. **A.** qPCR of cell-type dependent marker mRNAs. The differentiated cells are derived from three independent differentiation experiments. For statistical analysis, every group was measured in triplicate and differences were assessed using the one-sided statistical test. **B.** Immunofluorescence staining with cell type-specific markers. Nuclei are stained in blue using DAPI and the respective marker by antibody in red. Scale bar indicates 50µM length.

The astrocyte marker GFAP revealed a high mRNA and protein expression in the entire astrocyte population. In comparison to undifferentiated HNSC.100, GFAP was also moderately elevated in neurons and oligodendrocytes at the transcript level, where a few cells of the oligodendrocytes exhibit protein expression as well. Analysis of the neuronal marker MAP2 showed protein expression in every cell type, with the highest level in neurons. This observation correlates well with qPCR measurements showing the same trend (Fig. 3A, B). Immunofluorescence imaging of the oligodendrocyte marker CNPase selectively stained for oligodendrocytes (Fig. 3B and Supplementary Figure 4). However, on transcript level this marker did not seem to be suitable for the determination of differentiation states.

In summary, we validated that SOX2 antibody staining can be used as a neural stem cell marker, indicating the undifferentiated state of HNSC.100 cells. GFAP is suitable as an astrocyte marker by qPCR and antibody staining. Although GFAP was also reported to mark radial glia cells, it was absent in the NSC state and induced only upon differentiation. We therefore interpret its expression as indicative of astrocytic differentiation under our conditions. Furthermore, we observed in immunofluorescence staining that the neuronal marker MAP2 has limited diagnostic value for HNSC.100-derived neurons, and that CNPase is highly indicative as oligodendrocyte marker.

### Verification of differentiated subtypes

The brain is composed of many different subtypes of neurons and glial cells. In order to understand whether HNSC.100 cells can recapitulate some of those differentiation features, we assessed the differentiation potential beyond what was shown in Figures 1-3. For this purpose, we designed primer pairs for qPCR analyses of markers for immature (TUBB3), excitatory (TBR1), and inhibitory (GABA-B-R1) neurons, protoplasmic (S100β), fibrous (GFAP), or reactive (TIMP1) astrocytes, and progenitor (PDGFRA), immature (CNPase), or mature (PLP1) oligodendrocytes (Fig. 4A) [29–34]. The qPCR analysis of neurons shows a significant upregulation of the universal neuronal marker MAP2 and the excitatory neuronal marker TBR1. In contrast, the remaining two neuronal markers, TUBB3 (immature) and GABA-B R1 (inhibitory), decreased only moderately in comparison to undifferentiated HNSC.100 cells. Together, the marker analysis of the neuron population indicates a not-terminally differentiated state. Furthermore, the marker expression suggests differentiation directed towards excitatory neurons. The differentiated astrocytes show a strong elevation of GFAP levels and only very modest regulation of the S100β and TIMP1 markers, suggesting a homogenous culture of fibrous astrocytes. The marker analyses for oligodendrocytes indicated that we obtained a rather mature cell subtype, as on the one hand a significant increase of PLP1 and, on the other hand, no differences in PDGFRA and CNPase expression were observed.

**Fig. 4.**
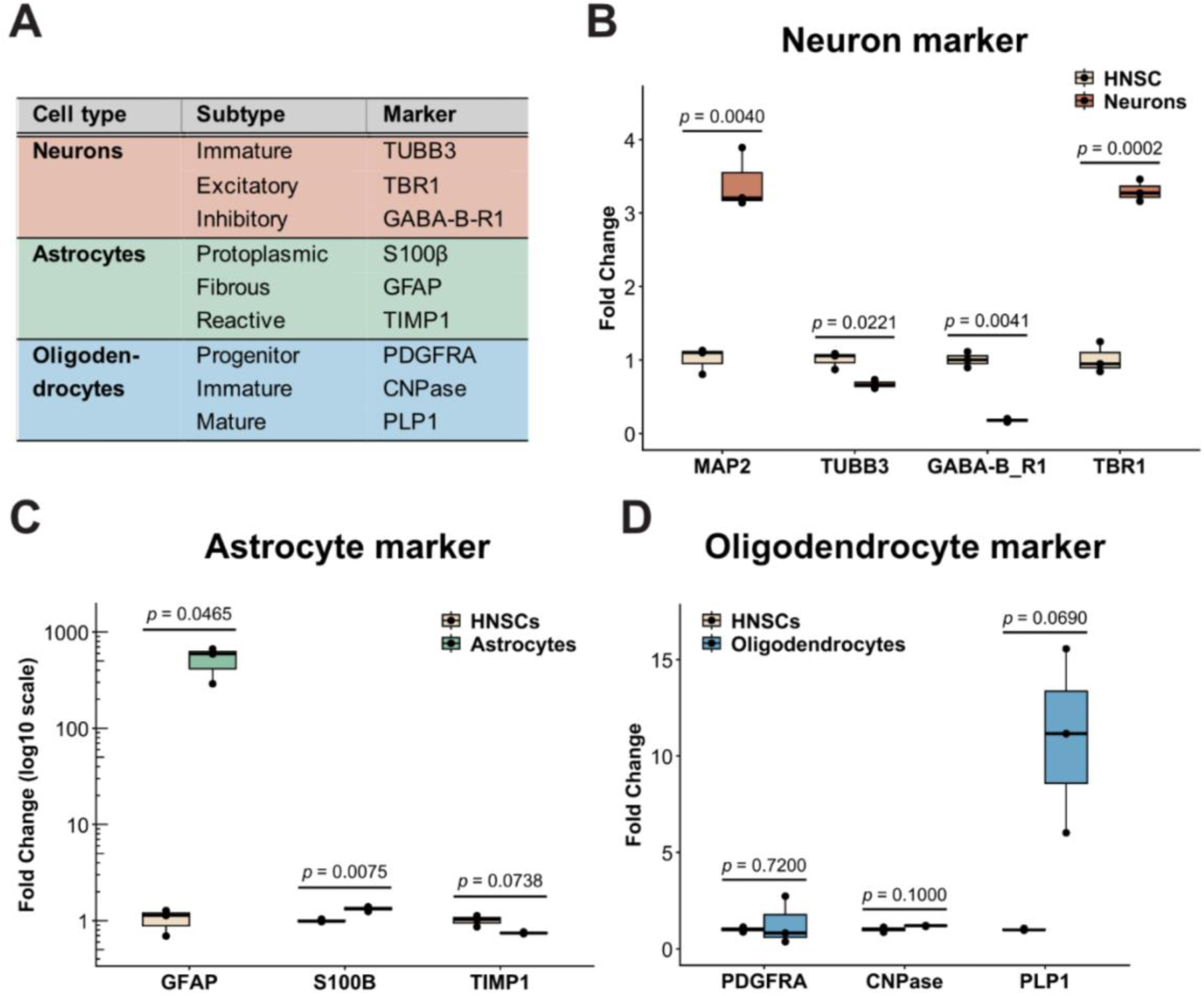
Subtype characterization of neurons, astrocytes and oligodendrocytes. Overview of specific subtype markers **(A)** and their expression levels in neurons **(B),** astrocytes **(C)** and oligodendrocytes derived from three independent differentiation experiments **(D).** Each experimental group was analyzed in three technical replicates, and statistical significance was determined using a two-sided statistical test.

The observed expression levels of these markers suggest that HNSC.100 cells successfully differentiate into fibrous astrocytes, whereas less prominent increase of mature-specific markers for neurons and oligodendrocytes indicates their less complete differentiation. Future work with additional markers could help to establish a finer-grained panel of cellular subtypes.

## Discussion

In this study, we assessed the human neural stem cell line 100 for its suitability as a model system for analyses of neural and neuronal functions. We provide an assessment of its differentiation potential alongside with a toolbox of validated differentiation markers.

A comparison of HNSC.100 with SH-SY5Y cells reveals that HNSC.100 offers a greater differentiation plasticity, potentially allowing for a wider range of studies. Karyotyping of both cell lines shows similar abnormalities (Supplementary Figure 1 and https://systemsbiology.uni.lu/shsy5y/). For instance, both cell lines have a very prominent duplication in the q-arm of chromosome 1. Other deviations from the wild-type heterozygous karyotype differ but are roughly comparable in their extent. It is often assumed that cancer-derived cell lines, such as SH-SY5Y, show greater chromosomal abnormalities than cell lines that were immortalized by engineering, such as HNSC.100 cells. Such a difference might be true for cell lines freshly generated by engineering. However, our analyses also suggest that proliferation of engineered immortalized cell lines inevitably results in similar abnormalities in their karyotypes as cancer cell lines. It should be noted, however, that these abnormalities in genotypes do not abolish the differentiation potential of these immortalized cell lines.

As a guideline, we recommend assessing the state of a gene of interest for a given project with the karyotyping result of this study to understand if it is present in wild-type copies or has adopted a genomic abnormality such as duplication or deletion. Together, these findings indicate that as for any immortalized cell line some limitations apply when using HNSC.100 cells, but that they are very well suited to study general principles of neur(on)al differentiation and function.

For a cell line to be suitable for a given project, one prerequisite is that the corresponding gene of interest is expressed in reasonable amounts. In general, several databases such as the GTEx Portal, Brain RNA-Seq, and The Human Protein Atlas serve as a great tool to assess the expression panel of a certain gene in their physiological states. However, these data usually do not include specific immortalized cell lines. To allow for an assessment of the suitability of the HNSC.100 cell line for a given project, we provide a list of all genes expressed above a certain threshold (Supplementary Table 3). Based on Illumina RNA sequencing data, we had set the minimal expression limit for inclusion in this list to 50 reads, as in our hands this turned out to be the lower limit for reliable quantification by qPCR. This list may not only serve as a reference to retrieve expression data for a gene of interest but could also serve as a reference point for the identification of additional differentiation markers. To prioritize disease-relevant genes, three databases for genes associated with neurodevelopment (DDG2P, SysNDD, DBD) and with neurodegeneration (DisGeNET, GWAS, PanelApp) were overlapped (Supplementary Table 4). The resulting list of common genes was filtered for high-confidence candidates, and their expression levels were analyzed in HNSC.100 cells. Most of the selected genes show substantial expression in HNSC.100 cells, supporting the suitability of this cell line as a model to study molecular mechanisms related to neurodevelopment and neurodegeneration (Fig. 5). Full lists of these disease-related genes are available in Supplementary Table 5 (neurodevelopmental disorders) and Supplementary Table 6 (neurodegenerative disorders).

**Fig. 5.**
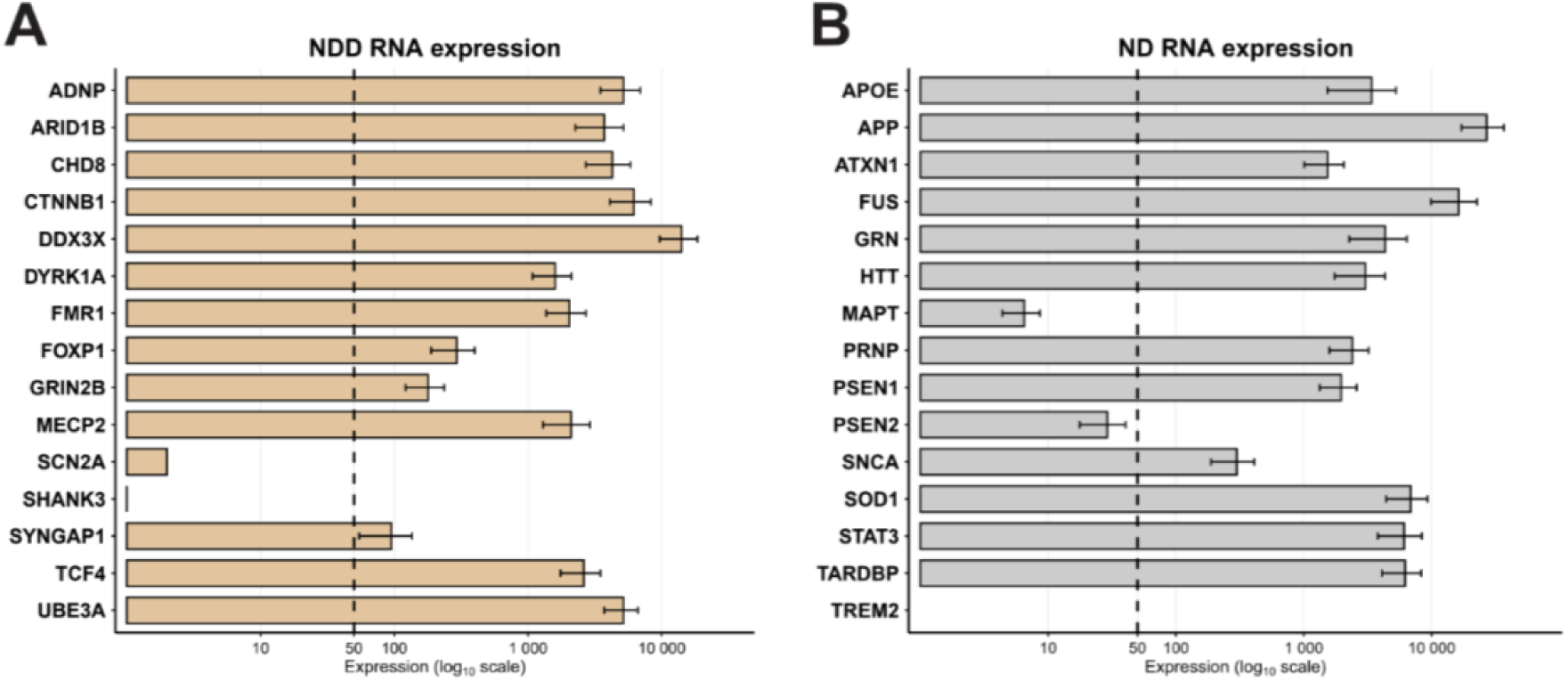
Expression panel of selected neural disease-related genes in HNSC.100 cells. Transcript counts of 15 selected genes associated with either neurodevelopmental diseases (NDDs) **(A)** or with neurodegenerative diseases (NDs) (**B**). Dashed vertical line indicates the threshold set in this study for reliable detection by qPCR. Genes expressed below this limit were excluded from Supplementary Table 3, but included in Supplementary Tables 5 and 6.

Together, this information shall provide the reader with an easy way to assess the suitability of HNSC.100 cells for their individual project.

The RNA-sequencing data were also used to compare the expression profiles of several marker genes in HNSC.100 with the immortalized cell lines ReNcell VM and ReNcell CX as well as human iPSC-derived NPCs (Supplementary Figure 3). This comparison yielded on one hand clear differences between these cell lines. On the other hand, the HNSC.100 line exhibited greater similarities to NPCs than both ReNcell lines in terms of the relative expression of NSC and proliferation markers compared to differentiation markers. Considering the overall low number of available immortalized neural stem-cell lines, this suggests that HNSC.100 provides a valuable expansion of the portfolio of available cell lines for orthogonal studies.

We demonstrated that HNSC.100 cells can be reliably differentiated into neurons, astrocytes, and oligodendrocytes under defined culture conditions, highlighting their robust differentiation capacity. The successful lineage specification was confirmed by distinct morphological characteristics and expression of lineage-specific markers such as MAP2 for neurons, GFAP for astrocytes, and CNPase for oligodendrocytes. Moreover, we established a comprehensive marker panel for subtype specification. This panel provides a practical framework that can be applied, adapted, and expanded by researchers. Nevertheless, additional work will be required to understand the full differentiation potential of the HNSC.100 cells. While in this study these cells fully differentiated into fibrous astrocytes after 16 days, a terminal differentiation of neurons and oligodendrocytes appeared to require cultivation for longer time periods than 28 and 35 days, respectively. Only with extended differentiation times will it become clear which exact neuronal and oligodendritic cell types result from the differentiation protocols of this study. After such validation of the terminally differentiated cells, attempts can be made to modify the differentiation paths into different but related fates.

Assessment of functional features, for instance, by electrophysiology could subsequently help to understand how well these cells recapitulate their physiological functions. Furthermore, several neural cell lines derived from glioblastoma and neuroblastoma have been used to establish 3D cell-culture models [35]. An obvious next step for HNSC.100 cells is therefore the exploration of their potential to self-organize into 3D cultures.

In summary, this study provides a detailed description of the human neural stem cell line HNSC.100 and offers a toolbox of specific markers to assess differentiation states and expression profiles of hundreds of disease-relevant genes. Our study might therefore serve as a reference point for using HNSC.100 cells to study the fundamental principles of neural function as well as related pathomechanisms.

## Materials and Methods

### Cell Culture

HNSC.100 cell line was kindly provided by R. Brack-Werner (Helmholtz Zentrum München) after a Material Transfer Agreement (ID: MEW-2379) had been signed with the Centro de Biologia Molecular “Severo Ochoa”, Universidad Autonoma de Madrid, Spain [12]. A basic description of this cell line, which is sometimes also referred to as hNS1 [14], is also available at https://www.cellosaurus.org/CVCL_B6QU. Cells were cultivated in Dulbecco’s Modified Eagle Medium/Nutrient Mixture F-12 (DMEM/F-12, 11320033, Gibco™) with the addition of 1% N-2 supplement (17502048, Gibco™), 1% BSA (P06-1403020, PAN Biotech), 1% Penicillin-Streptomycin (P06-07100, PAN Biotech), 0.5% HEPES buffer (P05-01100, PAN Biotech), 0.5% FBS (A5256801, Gibco™), 20 ng/ml EGF (in-house), and 20 ng/ml FGF-2 (in-house). Cells were split twice a week using Trypsin-EDTA (P10-025100, PAN Biotech) and reseeded in a T25 flask (83.3910.002, Sarstedt), which was beforehand coated for 30 min with 10% Poly-D-lysin (A3890401 diluted in DPBS (14190144, Gibco™) and washed twice with DPBS. Cells were stored in a cell incubator at 37 °C with 5% CO_2_. The media composition for HNSC.100 is listed in Supplementary Table 2.

### Differentiation

Differentiation into astrocytes was carried out for 16 days in previously coated 6-well plates for 30 min with 10% Poly-D-Lysin diluted in DPBS. To generate astrocytes, the aforementioned media (Cell Culture section) lacking EGF and FGF-2 was supplemented twice a week to the cells, as described elsewhere [12].

To generate neurons and oligodendrocytes, 6-well plate was first coated with 20 µg/ml poly-L-ornithine (A-004-C, Sigma-Aldrich) diluted in sterile H_2_O for 1h at 37 °C. This was followed by a second coating with 10 µg/ml laminin (23017015, Gibco™) diluted in sterile H_2_O for 2 h at 37 °C. Afterwards, plates were washed twice with DPBS.

Differentiation into neurons took 28 days was performed using a media composition recommended at the website of ThermoFisher [23]. In short, cells were supplemented twice a week with Neurobasal™ Medium (21103049, Gibco™) with the addition of 2% B-27™ supplement (17504044, Gibco™), 1% GlutaMAX™ supplement (35050061, Gibco™), and 1% Anti-Anti (15240062, Gibco™). Similarly, the oligodendrocyte differentiation lasted 28 days, where cells were supplemented with media containing the same components; however, additionally, 20 ng/ml PDGF-AA (in-house), and 30 ng/ml T3 (T5516, Sigma-Aldrich) were added. Notably, the used media composition was originally recommended for human embryonic stem cells (hESCs) [24] and optimized for HNSC.100 as part of this work.

Oligodendrocyte generation required an additional so-called pre-differentiation step of seven days prior to the 28-day-long differentiation procedure. For this purpose, cells were seeded on plates pre-coated with 10% Poly-D-lysin in DPBS for 30 min, followed by two DPBS wash steps. Cells were supplemented with Neurobasal™ Medium with the addition of B-27 (2%) and Gluta Max-1 (1%) supplements, Anti-Anti (1%), 20 ng/ml PDGF-AA (in-house), 10 ng/ml FGF (in-house), and 20 ng/ml NT3 (in-house). The media composition for HNSC.100 differentiation is listed in Supplementary Table 2.

### Real-Time quantitative PCR

RNA isolation was done using innuPREP RNA Mini Kit 2.0 (845-KS-2040010, iST) according to the manufacturer’s Protocol 2. The obtained RNA templates were used for reverse transcription using PrimeScript™ RT Master Mix (#RR036Q, Takara), resulting in cDNA with a final concentration 50 ng/µl. qPCR was performed using the TaqMan probe approach and innuMIX qPCR MasterMix (845-AS-1200015, iST) containing innuTaq Hot-A DNA Polymerase. Each sample was performed in triplicate, where each cell type was independently generated three times for statistical analysis. Samples were added to a 96-well plate (Sarstedt) using low-adherence nuclease-free tips (Nerbe) and sealed with a transparent sealing foil (Sarstedt). Plates were centrifuged at 200 × g for 1 min and transferred to the qTower^3^ (Analytik Jena) machine. Each reaction was performed for the gene of interest and two housekeeping genes: *GAPDH* and *RPL32* for normalization using the ΔΔC method [36]. Statistical differences between groups were validated using one-sided and two-sided Student’s t-tests or a U Mann-Whitney test for independent samples, depending on the Shapiro test results. One-sided statistical tests were applied where predefined directional hypotheses were formulated such as expected upregulation of lineage-specific differentiation markers or downregulation of NSC markers upon differentiation. Two-sided tests were used for analyses where no predefined direction of regulation was assumed, such as subtype specification. The list of used oligonucleotides for qPCR is available in Supplementary Table 7, where each gene-of-interest design was validated by testing the primer efficiency. For this purpose, a new target was tested in a series of five serial dilutions, beginning with an undiluted sample. Subsequent dilutions were prepared by a 10-fold dilution at each step. Then, the measured cycle (Ct) values for each dilution was plotted against the logarithm of the dilution factors, resulting in a generated amplification curve, and the calculated curve slope [37].

### Immunostaining and fluorescence microscopy

For protein expression analysis, un/-differentiated cells were cultivated on sterile and coated glass coverslips as described in the Differentiation section. Samples were washed with DPBS and fixed with 3.7% Formaldehyde (F8775-25ML, Sigma-Aldrich) in DPBS for 7-10 min. Afterward, cells were washed two times with DPBS, followed by membrane permeabilization using 0.5% Triton X-100 (X100, Sigma-Aldrich) in PBS for 5 min at room temperature and two washes with DPBS. Next, slides were placed on parafilm and blocked for 30 min with a blocking solution composed of 1% Goat serum (16210064, Gibco™) and 0.05% Tween-20 (A4974.0500, AppliChem) in PBS. Then, cells kept on parafilm were incubated overnight with the primary antibody diluted in a blocking solution at 4 °C. To prevent drying of samples, a parafilm slice with samples was placed in a box filled with water-soaked paper tissues and covered with a lid. The next day, slides were washed with PBS three times for 5 min, followed by 1 h incubation with secondary antibody in blocking solution in the dark at room temperature. Slides were again washed with PBS three times for 5 min and stained with DAPI (D1306, Thermo Scientific™) at a final concentration of 0.5 µg/ml. Finally, cells were washed twice with PBS and embedded in a beaker with water. Then, the slides were dried with a paper tissue and mounted onto a microscopic slide using ProLong Diamond Antifade Mountant (P36961, Invitrogen™). Slides were left for drying at room temperature in the dark for at least 5 h and stored in the fridge. The sample imaging was taken using a Leica DMi8 microscope and Leica Imaging software. Antibodies used for the IF analysis are listed in Supplementary Table 8.

### RNA sequencing

For validation of transcript expression in HNSC.100 cells, Illumina sequencing after reverse transcription was performed. For this purpose, cells were independently cultivated in quadruplicates and RNA isolation was performed using innuPREP RNA Mini Kit 2.0 for 1 Mio of cells as an input. RNA concentration was measured using Qubit™ RNA BR Assay Kit (Q10210, Invitrogen™) and the QFX Fluoremeter (DeNovix). Additionally, the quality of RNA was determined using High Sensitivity RNA ScreenTape (5067-5579, Agilent) for the TapeStation (Agilent). Next, 1 µg of total RNA per replicate was used for mRNA isolation using NEB Next Poly(A) mRNA Magnetic Isolation Module (E7490S, NEB) according to the manufacturer’s instructions. The obtained sample was then used for the cDNA generation and NGS library preparation using NEBNext^®^ Ultra^™^ II Directional RNA Library Prep Kit for Illumina^®^ (E7765, NEB) following the manufacturer’s guidance. Generated libraries were validated using High Sensitivity D1000 Screen Tape (5067-5584, Agilent), and sequenced at the Gene Center, LMU Munich using a 60 bp paired-end setting for 30-35 million reads per replicate. The data can be accessed via GEO accession number: GSE317784.

### Comparison of immortalized and iPSC-derived neural stem cells

Publicly available bulk RNA-seq datasets from the immortalized neural stem cell lines ReNcell CX and ReNcell VM (GEO accession numbers: GSE92839, GSE89623) [18,19] and iPSC-derived NPCs [20] were analyzed. All available biological replicates were included. For comparability with GEO-provided TPM expression matrices, TPM values for HNSC.100 and iPSC-derived NPCs were calculated using gene lengths provided in the NCBI annotation (GRCh38.p13). Expression of lineage-specific marker genes was visualized as heatmaps using log10(TPM + 1) transformed values.

### Karyotyping

For karyotype analysis genomic DNA (gDNA) was isolated using the DNeasy Blood & Tissue Kit (Qiagen, #69504). The array-based karyotyping was conducted by Life & Brain GmbH (Bonn, Germany) on HNSC.100 wild-type gDNA using the Illumina BeadChip platform. Raw data were processed and analyzed with GenomeStudio v2.0 (Illumina).

### Lipotransfection and nucleofection of HNSC.100

HNSC.100 were seeded to achieve around 80% confluency at the time of transfection. The lipotransfection was performed using the Lipofectamine 3000 transfection kit (L3000, Invitrogen™) according to the manufacturer instructions. The P3 Primary Cell 4D-Nucleofector® X Kit and the Amaxa 4D device (Lonza) were used to electroporate the cells according to the manufacturer instructions. For assessing transfection, 2 µg of the control vector pmaxGFP^TM^ Vector (Lonza) was used for each transfection. The cells were incubated for three days after transfection and GFP signal was analyzed using the Countess 3 FL (Thermo Fisher) automated cell counter with an EVOS™ light cube GFP 2.0.

### Identification of genes associated with neurodevelopmental and neurodegenerative diseases by database overlaps

Gene-level evidence for neurodevelopmental diseases (NDDs) was integrated from DDG2P, SysNDD, and the DBD database. Evidence for neurodegenerative diseases (NDs) was collected from PanelApp, DisGeNET, and NHGRI-EBI GWAS Catalog. The exact disease terms, web links, and dataset versions used for filtering are provided in Supplementary Table 4. All gene annotations were harmonized and merged into a unified evidence scoring (High, Moderate, and Low) based on the criteria summarized in Supplementary Table 4. For each gene, the number and strength of supporting sources were aggregated to derive a combined evidence level. The number of High (H), Moderate (M), and Low (L) ratings across all sources was counted and genes were classified as Very High if H = 3; High if H ≥ 2; Moderate if (H ≥ 1 and M ≥ 1) or M ≥ 2; and Low if L ≥ 2 or if evidence was available from only one source.

### Generation of schematic drawings

Cartoons for figures were generated using Biorender (license number IRICKYOB).

## Acknowledgement

We are grateful to Vera Roman, Manuel Stech Domene and Silvia Marcato for their contribution and would like to thank Helmut Blum and Stefan Krebs for Illumina sequencing. This work was supported by the Deutsche Forschungsgemeinschaft (NI 2094/11-1, project number: 541627224 to DN), by the PURA Syndrome Foundation and by PURA Syndrome Deutschland e.V..

## Supplementary Data

**Supplementary Figure 1.**
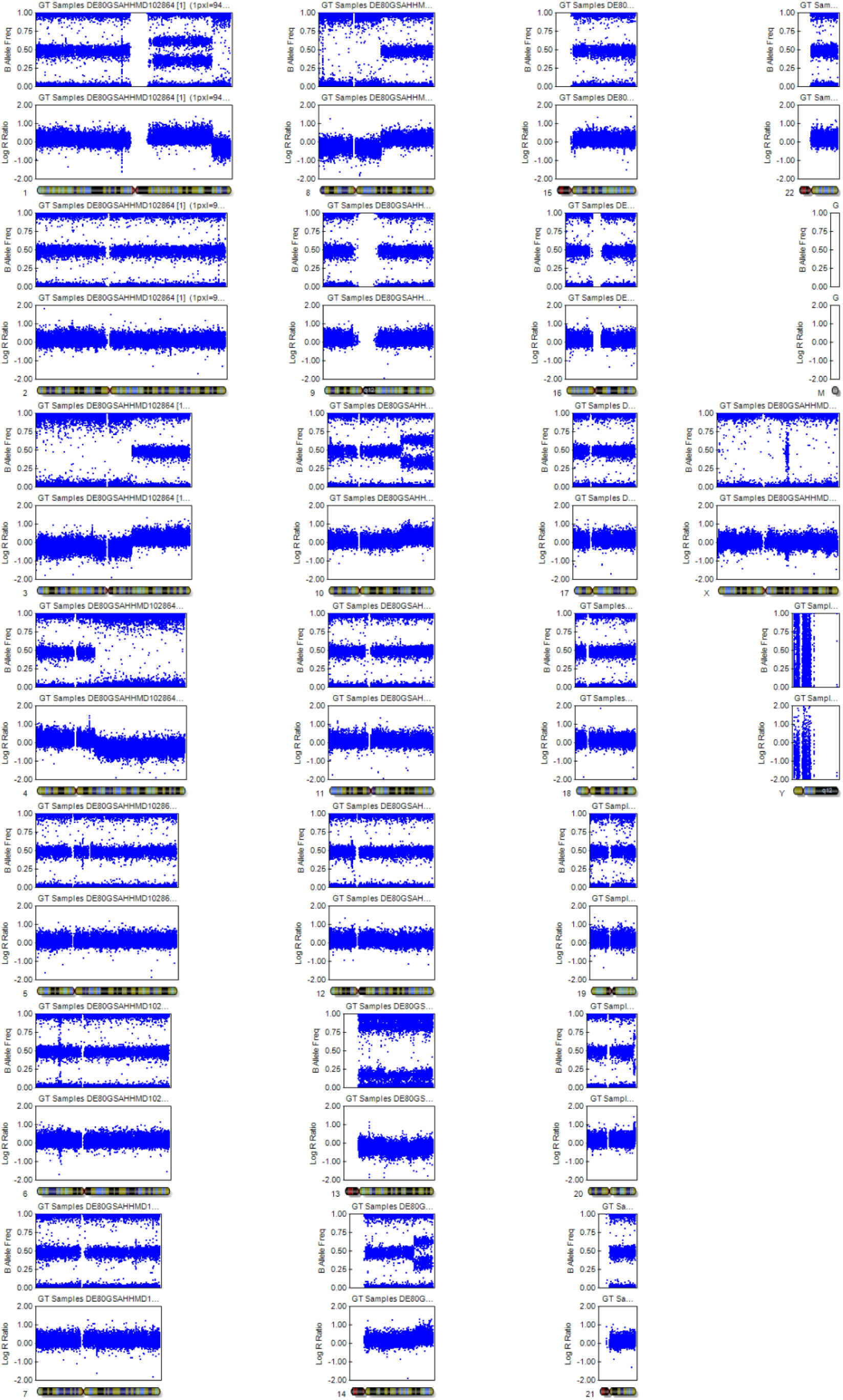
Karyotype of HNSC.100 cells.

**Supplementary Figure 2.**
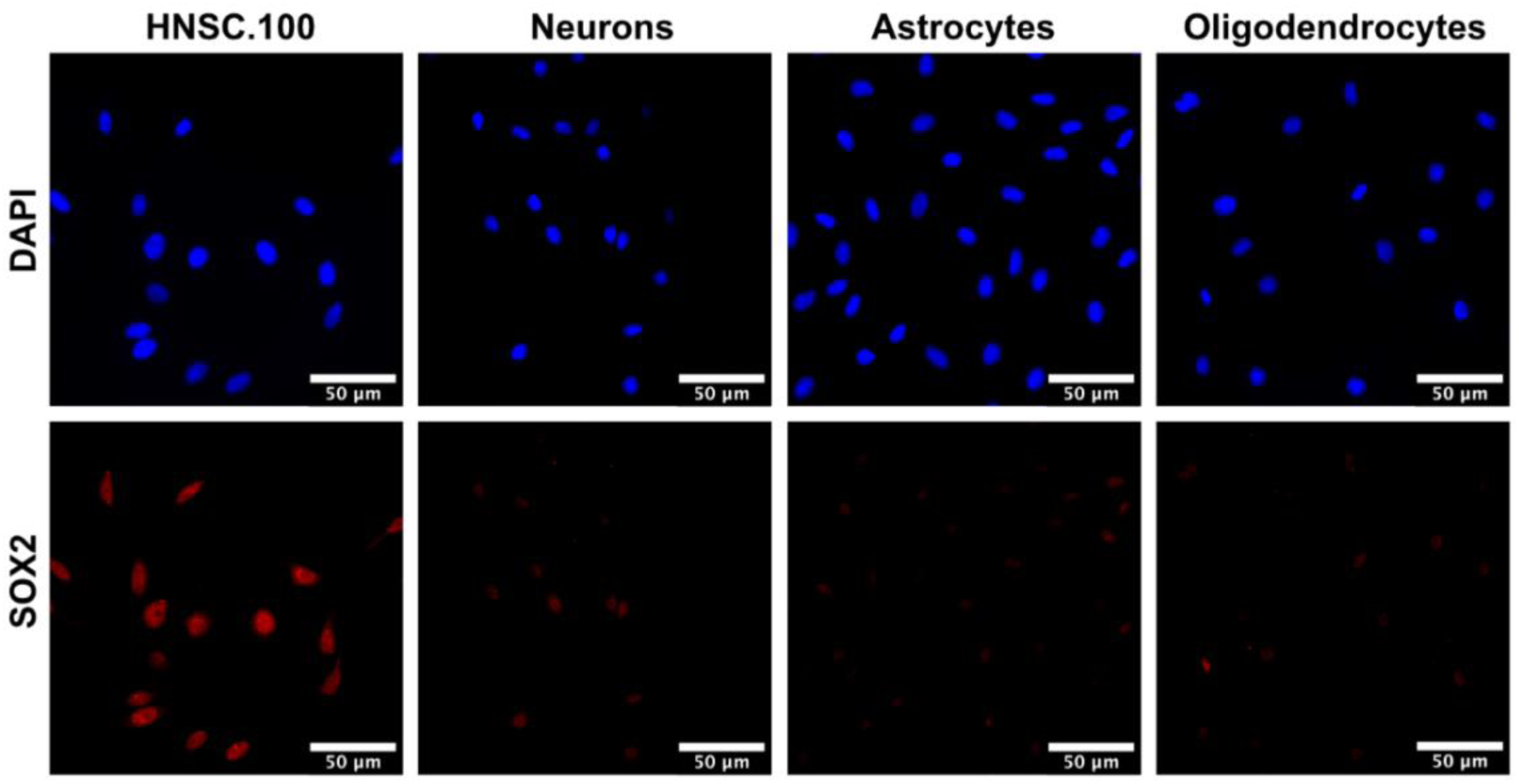
Single-channel visualization of SOX2 staining in undifferentiated and differentiated HNSC.100 cells. Representative immunofluorescence images showing SOX2 (red) and DAPI (blue) channels separately for undifferentiated HNSC.100 cells, neurons, astrocytes and oligodendrocytes.

**Supplementary Figure 3.**
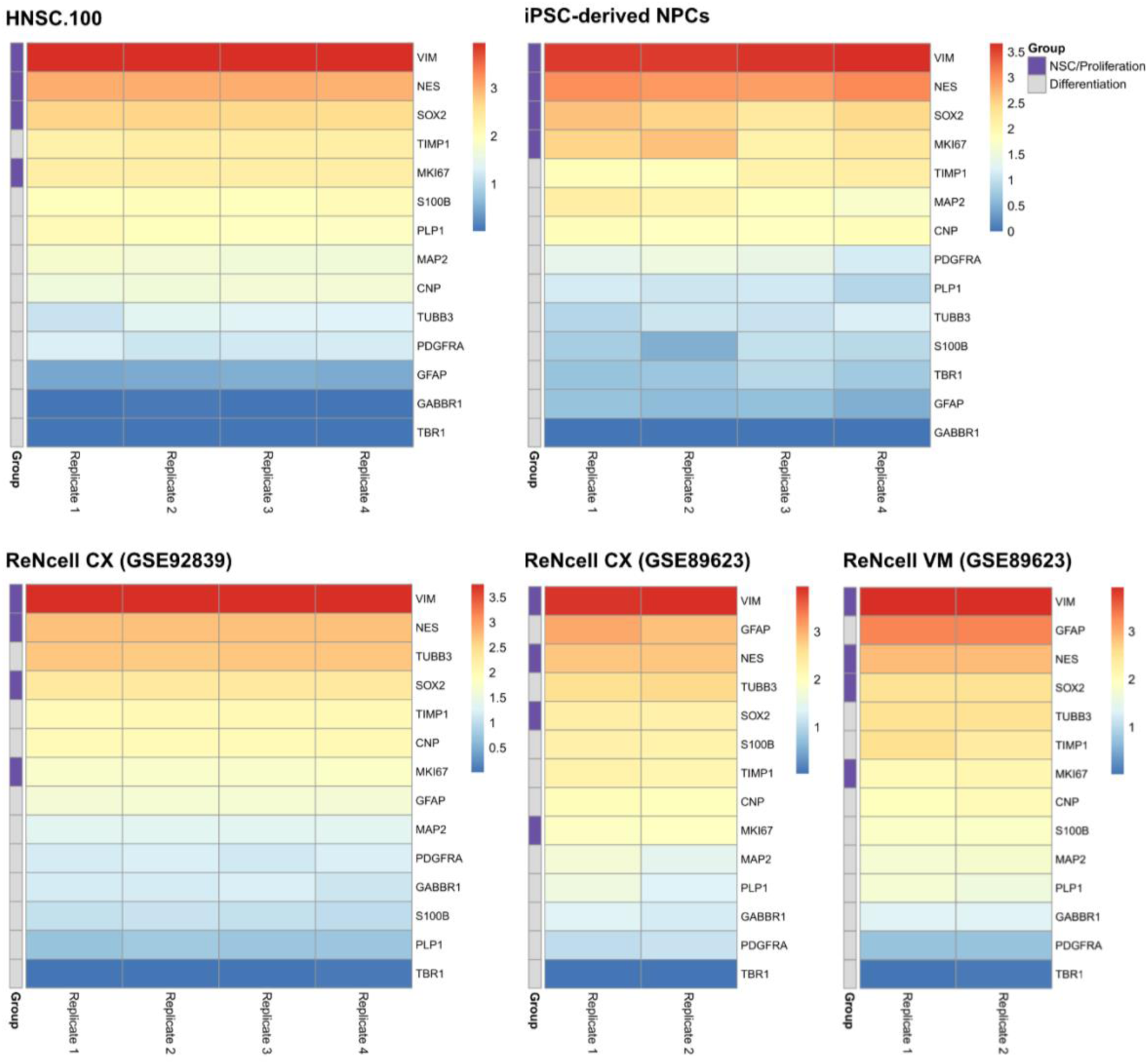
Comparison of lineage marker expression across neural stem cell models. Heatmaps show relative expression of neural stem cell/ proliferation markers (purple) and differentiation markers (grey) representing neuronal, astrocytic, and oligodendrocytic lineages. Data include RNA sequencing from HNSC.100 and publicly available RNA sequencing datasets from the immortalized neural stem cell lines ReNcell CX and VM, as well as iPSC-derived neural progenitor cells (NPCs). The two different RenCell CX heatmaps correspond to independent datasets (GEO accession numbers indicated). Expression values are shown as log10(TPM + 1).

**Supplementary Figure 4.**
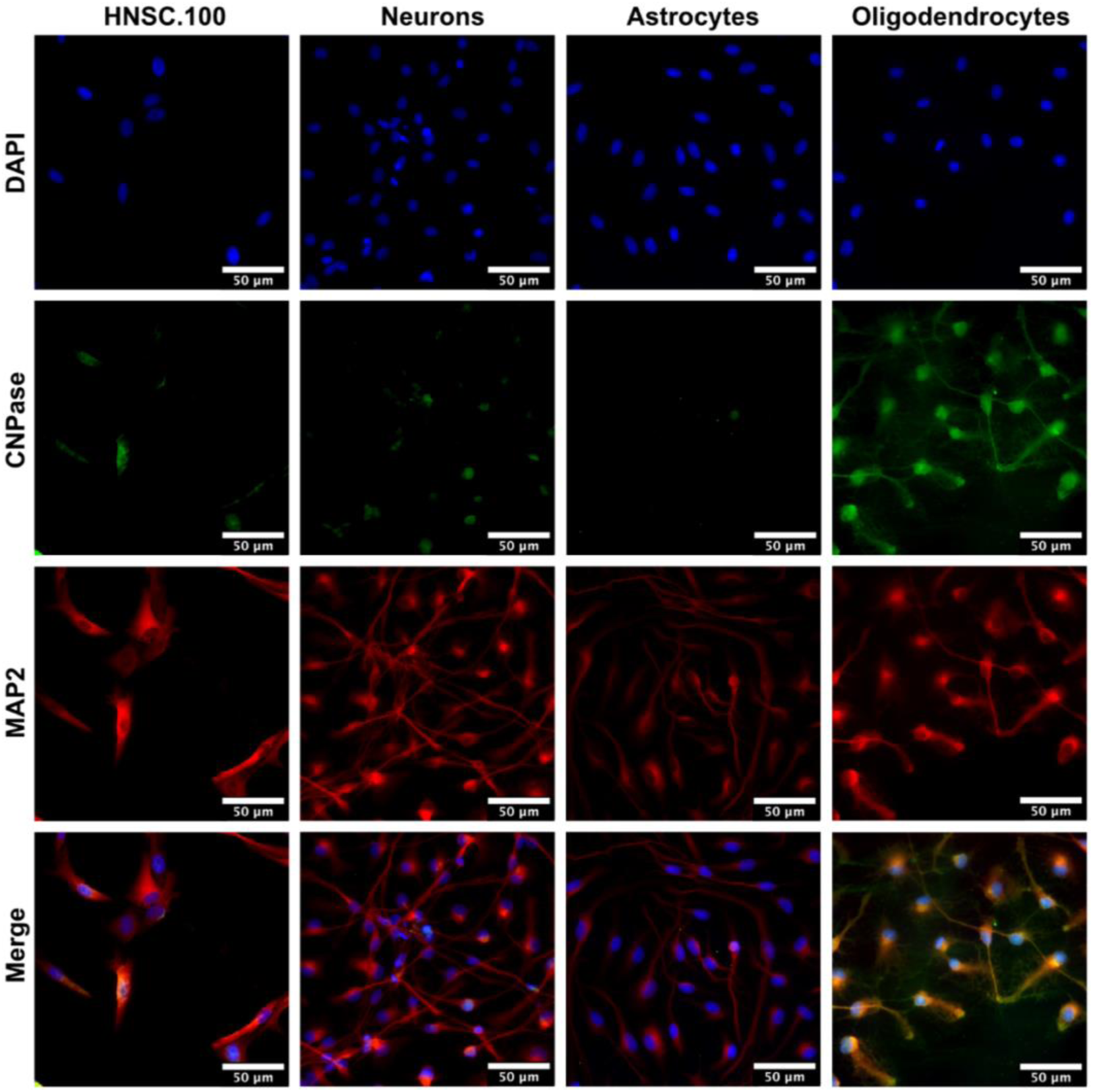
Co-staining of MAP2 and CNPase in undifferentiated and differentiated HNSC.100 cells. Representative immunofluorescence images showing co-staining of the neuronal marker MAP2 (red), the oligodendrocyte marker CNPase (green), and DAPI (blue) in undifferentiated HNSC.100 cells and cells differentiated into neurons, astrocytes, and oligodendrocytes.

**Supplementary Table 1.** Tabular representation of major chromosomal abnormalities from Supplementary Figure 1:

| chromosome | region | type | copy number | ISCN |
| --- | --- | --- | --- | --- |
| 3 | p26.3–q21.2 | deletion | ×1 | del(3)(p26.3q21.2) |
| 4 | q21.1–q35.2 | deletion | ×1 | del(4)(q21.1q35.2) |
| 8 | p23.3–q21.12 | deletion | ×1 | del(8)(p23.3q21.12) |
| 10 | q23.33–q26.3 | duplication | ×3 | dup(10)(q23.33q26.3) |
| 13 | q12.11–q34 | deletion | ×1 | del(13)(q12.11q34) |
| 14 | q31.2–q32.33 | duplication | ×3 | dup(14)(q31.2q32.33) |
| X | complete | deletion | ×1 | -X |

**Supplementary Table 2.**
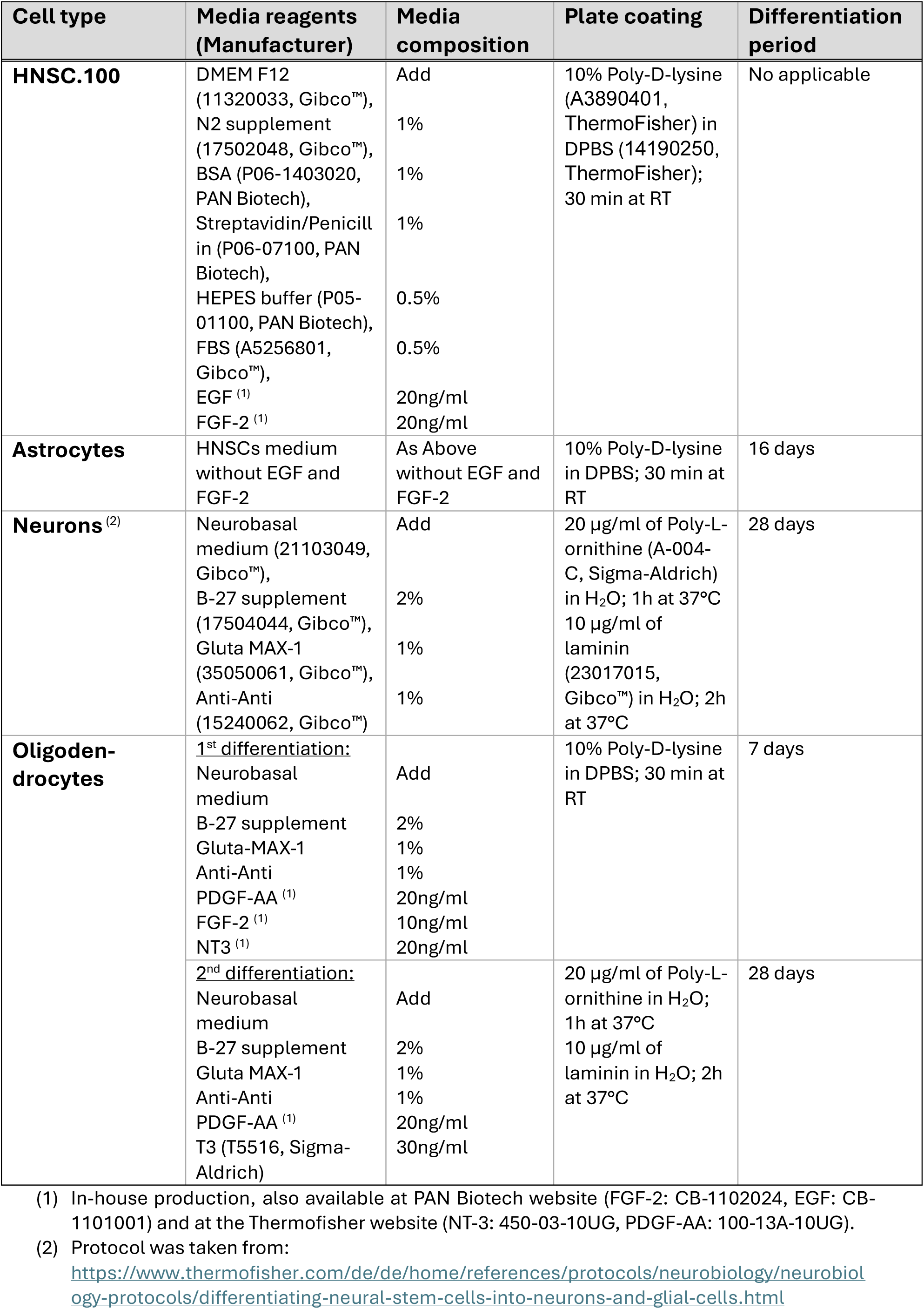
Conditions for cultivation and differentiation of HNSCs.

| Cell type | Media reagents (Manufacturer) | Media composition | Plate coating | Differentiation period |
| --- | --- | --- | --- | --- |
| **HNSC.100** | DMEM F12 (11320033, Gibco™), N2 supplement (17502048, Gibco™), BSA (P06-1403020, PAN Biotech), Streptavidin/Penicill in (P06-07100, PAN Biotech), HEPES buffer (P05-01100, PAN Biotech), FBS (A5256801, Gibco™), EGF <sup>(1)</sup> FGF-2 <sup>(1)</sup> | Add<br>1%<br>1%<br>1%<br>0.5%<br>0.5%<br>20ng/ml<br>20ng/ml | 10% Poly-D-lysine (A3890401, ThermoFisher) in DPBS (14190250, ThermoFisher); 30 min at RT | No applicable |
| **Astrocytes** | HNSCs medium without EGF and FGF-2 | As Above without EGF and FGF-2 | 10% Poly-D-lysine in DPBS; 30 min at RT | 16 days |
| **Neurons** <sup>(2)</sup> | Neurobasal medium (21103049, Gibco™), B-27 supplement (17504044, Gibco™), Gluta MAX-1 (35050061, Gibco™), Anti-Anti (15240062, Gibco™) | Add<br>2%<br>1%<br>1% | 20 µg/ml of Poly-L-ornithine (A-004-C, Sigma-Aldrich) in H<sub>2</sub>O; 1h at 37°C<br>10 µg/ml of laminin (23017015, Gibco™) in H<sub>2</sub>O; 2h at 37°C | 28 days |
| **Oligodendrocytes** | <u>1<sup>st</sup> differentiation:</u><br>Neurobasal medium<br>B-27 supplement<br>Gluta-MAX-1<br>Anti-Anti<br>PDGF-AA <sup>(1)</sup><br>FGF-2 <sup>(1)</sup><br>NT3 <sup>(1)</sup> | Add<br>2%<br>1%<br>1%<br>20ng/ml<br>10ng/ml<br>20ng/ml | 10% Poly-D-lysine in DPBS; 30 min at RT | 7 days |
|  | <u>2<sup>nd</sup> differentiation:</u><br>Neurobasal medium<br>B-27 supplement<br>Gluta MAX-1<br>Anti-Anti<br>PDGF-AA <sup>(1)</sup><br>T3 (T5516, Sigma-Aldrich) | Add<br>2%<br>1%<br>1%<br>20ng/ml<br>30ng/ml | 20 µg/ml of Poly-L-ornithine in H<sub>2</sub>O; 1h at 37°C<br>10 µg/ml of laminin in H<sub>2</sub>O; 2h at 37°C | 28 days |

(1) In-house production, also available at PAN Biotech website (FGF-2: CB-1102024, EGF: CB-1101001) and at the ThermoFisher website (NT-3: 450-03-10UG, PDGF-AA: 100-13A-10UG).
(2) Protocol was taken from:
https://www.thermoFisher.com/de/de/home/references/protocols/neurobiology/neurobiology-protocols/differentiating-neural-stem-cells-into-neurons-and-glial-cells.html

**Supplementary Table 3:** See Excel file Supplementary_Table_3.xlsx

**Supplementary Table 4.** Search and scoring criteria for each database with respect to neurodevelopment (DDG2P, SysNDD, DBD) and neurodegeneration (PanelApp, DisGeNET, and NHGRI-EBI GWAS Catalog). Database entries were derived from:

| Database | Search Criteria | Rule | Evidence level |
| --- | --- | --- | --- |
| **DDG2P** | All DDG2P entries labeled as “developmental disorder (DD)” | Confidence = “Definitive” | High |
|  |  | Confidence = “Strong”, “Probable” | Moderate |
|  |  | Confidence = “Limited”, “Possible” or lower | Low |
| **SysNDD** | All curated monogenic NDD genes (entities_category annotated) | Category = “Definitive” and NDD phenotype = TRUE | High |
|  |  | Category = “Strong” or “Moderate” | Moderate |
|  |  | All others | Low |
| **DBD** | All genes with Tier 1–4 classification (de novo / inherited mutation evidence) | Tier = 1 | High |
|  |  | Tier = 2 or 3 | Moderate |
|  |  | Tier = 4 or AR (autosomal recessive only) | Low |
| **PanelApp** | Adult onset neurodegenerative disorder | Panel rating = Green | High |
|  |  | Panel rating = Amber | Moderate |
|  |  | Panel rating = Red | Low |
| **DisGeNET** | Major neurodegenerative disorders: Alzheimer’s, Parkinson’s, ALS/FTD, Huntington’s, spinocerebellar ataxias, prion diseases, dementia | Score $\geq 0.6$ | High |
| | | $0.3 \leq \text{Score} < 0.6$ | Moderate |
| | | Score $< 0.3$ | Low |
| **GWAS Catalog (NHGRI-EBI)** | Filtered for DISEASE/TRAIT terms: “Alzheimer”, “Parkinson”, “amyotrophic lateral sclerosis”, “frontotemporal”, “Huntington”, “prion”, “ataxia”, “dementia”, “neurodegenerative” | $P \leq 5 \times 10^{-8}$ | High |
| | | $5 \times 10^{-8} < P \leq 1 \times 10^{-5}$ | Moderate |
| | | $P > 1 \times 10^{-5}$ | Low |

- G2P: https://www.ebi.ac.uk/gene2phenotype/
- SysNDD: https://sysndd.dbmr.unibe.ch/
- DBD: https://dbd.geisingeradmi.org/
- PanelApp: https://panelapp.genomicsengland.co.uk/
- DisGeNET: https://disgenet.com/
- NHGRI-EBI GWAS Catalog: https://www.ebi.ac.uk/gwas/

**Supplementary Table 5:** See Excel file Supplementary_Table_5.xlsx

**Supplementary Table 6:** See Excel file Supplementary_Table_6.xlsx

**Supplementary Table 7.**
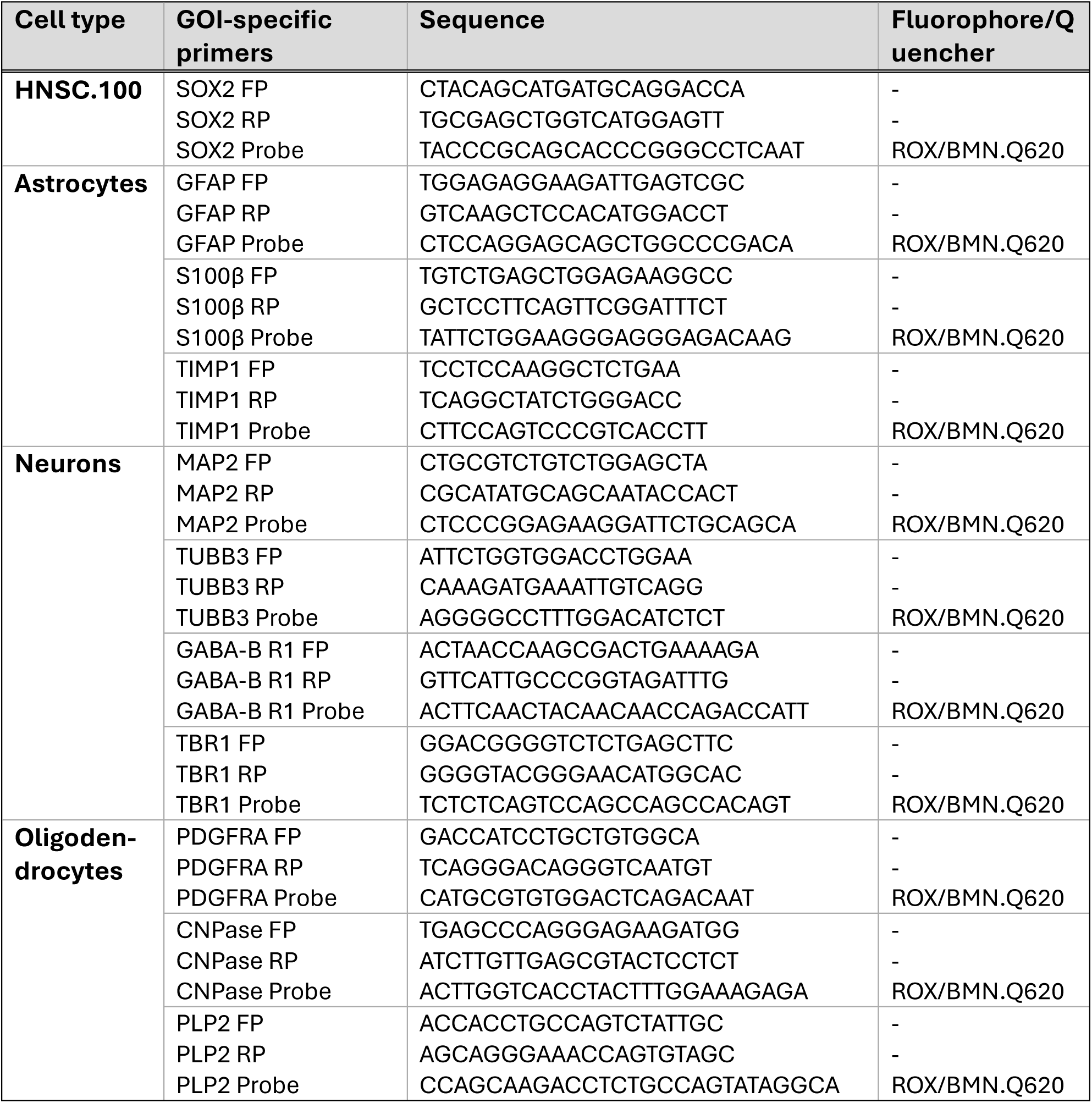
qPCR setups for cell-type validation.

**Supplementary Table 8.** Antibodies used for cell-type validation.

| Cell type | Targeted protein (host) | Dilution | Supplier | Cat. number |
| --- | --- | --- | --- | --- |
| **HNSC.100** | SOX2 (rabbit) | 1:500 | Proteintech | 11064-1-AP |
| **Astrocytes** | GFAP (rabbit) | 1:500 | Proteintech | 16825-1-AP |
| **Neurons** | MAP2 (rabbit) | 1:500 | Proteintech | 17490-1-AP |
| **Oligodendrocytes** | CNPase (mouse) | 1:200 | Proteintech | 66729-1-Ig |

**Supplementary Table 9.**
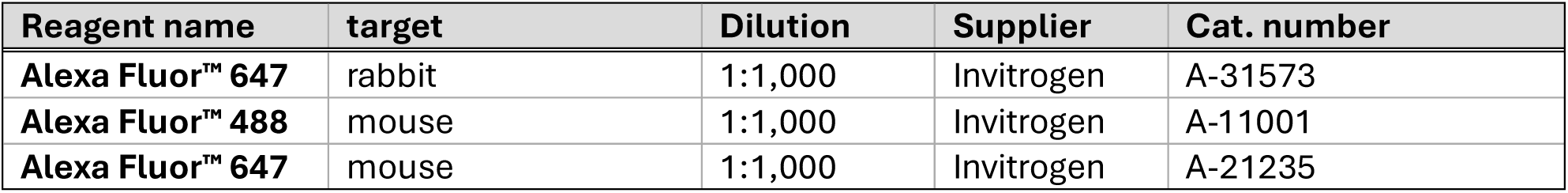
Secondary antibodies used for cell-type validation.

## Notes

### Competing Interest Statement

The authors have declared no competing interest.

### Summary of Updates

Correction of formatting issues and few errors. Clarification of differentation protocol (e.g. in Fig. 2 and Supplementary Table 2). Statistics were reanalysed and in some instances adapted (e.g. Figure 4). New figures with additional data were added (Supplementary Figure 2-4). New Supplementary Table with list of antibodies was added.

